# Investigation of expression of genes encoding glycogen degrading enzymes in *Gardnerella swidsinskii* and identification of reference genes for quantitative real-time PCR

**DOI:** 10.1101/2024.03.02.583113

**Authors:** Andy Kim, Champika Fernando, Divanthika Kularatne, Janet E. Hill

## Abstract

*Gardnerella* spp. express and export enzymes for the breakdown of glycogen into glucose, maltose, and malto-oligosaccharides for consumption by the vaginal microbiota but how the expression of these “public goods” is affected by substrate and product levels in the environment is not known. Accurate measurement of relative gene expression using real-time quantitative PCR relies on the identification of appropriate reference genes whose expression levels remain constant under the conditions of the study. Currently, no reference genes have been identified for gene expression analysis of *Gardnerella* spp. The objectives of this study were to identify reference genes and apply them in determining the relative gene expression levels of genes encoding α-amylase and α-amylase-pullulanase in media supplemented with substrate (glycogen) or a preferred product (maltotriose). Ten candidate reference genes were evaluated and analysis of Cq values from qPCR using multiple algorithms identified *uppS* (encoding polyprenyl diphosphate synthase) as the top comprehensively ranked reference gene followed by *gatA* (encoding Asp-tRNA/Glu-tRNA amidotransferase subunit gatA). Interpretation of the Cq values for α-amylase and α-amylase-pullulanase was performed by applying these two reference genes in the calculation of relative gene expression levels. α-amylase-pullulanase gene expression was upregulated in media supplemented with 1% glycogen in comparison to media supplemented with 1% maltotriose suggesting a regulatory mechanism in *G. swidsinskii* that responds to nutrient availability. No significant difference in gene expression of α-amylase was observed suggesting expression is not influenced by substrate availability. The RNA purification protocol and reference genes validated in this study will be useful in future studies of gene expression in *Gardnerella*.

**Importance:** Knowledge of the factors affecting growth of vaginal microbiota is critical to understanding how vaginal dysbiosis is initiated and maintained. Overgrowth of *Gardnerella* species including *G. swidsinskii* is a hallmark of bacterial vaginosis. These organisms break down vaginal glycogen and the products become available for uptake by *Gardnerella* and other microbiota. Measuring how expression of genes encoding glycogen degrading enzymes relates to relative abundance of substrate and products in the environment requires development of protocols for RNA purification and identification of reference genes for RT-qPCR.

## Introduction

Abundant growth of *Gardnerella* spp. (including *G. vaginalis, G. swidsinskii, G. leopoldii*, and *G. piotii*) in the vaginal microbiome is a hallmark of bacterial vaginosis (BV), a condition in which normally dominant lactobacilli are replaced by an overgrowth of mixed facultative and anaerobic species (1). Although multiple species may coexist the vaginal microbiomes of women with or without BV, the relative abundance of the species varies among individuals (2). *G. vaginalis* and *G. swidsinskii* are frequently the most abundant of the *Gardnerella* species; they co-occur often, and their abundance is significantly associated with clinical signs of odour and vaginal discharge (2). Given the phenotypic differences among *Gardnerella* species (3) their hypothesized role in initiating BV (4) and reports of differential associations with clinical signs of BV, understanding the determinants of *Gardnerella* spp. population structure in the vaginal microbiome is important. Previous investigations have shown that competition for nutrients is a contributor to determining *Gardnerella* population structure (5).

Within the vaginal microbiome, glycogen is an important carbon source competed for by the diverse microbiota (6). Glycogen is hydrolyzed by human and/or bacterial enzymes into glucose, maltose and malto-oligosaccharides, which are then able to be utilized by the vaginal microbiota (7, 8). We have recently demonstrated that all *Gardnerella* spp. produce extracellular α-amylase and α-amylase-pullulanase that can release maltose and malto-oligosaccharides from glycogen (9) and that *Gardnerella* spp. show more growth in media supplemented with maltotriose and maltotetraose relative to glucose or maltose (10). The glycogen degrading enzymes produced by *Gardnerella* spp. comprise “public goods” for the vaginal microbiota and it is not known if they are constitutively expressed or if their expression is affected by the relative levels of products (malto-oligosaccharides) or substrate (glycogen). Production and export of glycogen degrading enzymes (including the >100 kDa α-amylase-pullulanase) poses an expense to the bacteria and so regulation of gene expression in response to substrate and product levels could be beneficial. Knowledge of gene expression in *Gardnerella* spp. is sparse and limited by the current lack of reference genes for transcript quantification.

Reverse transcriptase quantitative real-time polymerase chain reaction (RT-qPCR) is a powerful method for targeted quantification of gene expression (11). Reliable RT-qPCR results depend on control of several variables including the amount of starting material, reverse-transcriptase and polymerase enzyme efficiencies, and differences among cells in overall transcriptional activity (11). Differences in quality and integrity of RNA samples, efficiency of cDNA synthesis and PCR efficiency can drastically influence signal normalization (12, 13). In an effort to reduce the variability caused by these factors, the MIQE guidelines were established to provide directions for researchers in developing qPCR experiments (14). RNA integrity and quality is one of the many standards to be set by these guidelines as it has been shown that failure to account for individual RNA integrity can lead to false conclusions and publication retraction (15). High quality RNA is quantified by an RNA Integrity Number (RIN) ranging from 1 to 10 (1 = completely degraded RNA, 10 = most intact RNA) with a RIN of 8 or higher representing high quality RNA, is a prerequisite for reliable quantification of transcription levels and sequencing (16–19). RT-qPCR generally involves the normalization of expression levels of the genes of interest to the expression levels of suitable reference genes (20). It has been shown that the expression levels of many commonly used reference genes differ drastically under different treatment conditions (21, 22). Therefore, it is necessary to conduct reference gene identification on a case-by-case basis, even for the same species, as there is no universal gene that can be used as an internal control for all applicable studies (20).

Here we developed a protocol for purifying high quality RNA from *Gardnerella* spp., identified suitable reference genes for the conditions of our study, and applied these in quantification of α-amylase and α-amylase-pullulanase gene expression in *G. swidsinskii* grown in media supplemented with glycogen (substrate) or maltotriose (products).

## Materials and Methods

### Bacterial strains

*Gardnerella swidsinskii* strains NR016, NR020, and NR021 were cultured on Columbia sheep blood agar plates for 48 hours at 37°C under anaerobic conditions (GasPak™ EZ Anaerobe Gas Generating Pouch System with Indicator, BD) from isolates stored at -80°C. Modified NYC III medium (mNYC III) was prepared, supplemented with 1% filter sterilized oyster glycogen (9005-79-2, Sigma-Aldrich), maltotriose (J66491.03, ThermoFisher), or D-glucose (D16-500, FisherChemical). mNYC III is NYC III (ATCC Medium 1685) with 10% (v/v) heat inactivated fetal bovine serum instead of horse serum and no additional glucose (9). Colonies isolated from growth on Columbia sheep blood agar plates were used to inoculate mNYC III media and cultures were incubated for 48 hours at 37°C under anaerobic conditions (GasPak™ EZ Anaerobe Container System Sachets, BD).

### Growth curves

To determine which OD_600_ readings corresponded to exponential phase, a 48-hour broth culture (mNYC III supplemented with 1% oyster glycogen or maltotriose) was diluted 1:10 and 200 µL aliquots of the diluted sample were pipetted into 8 wells of a flat bottom 96-well plate (VWR Tissue Culture Plate, Non-treated, Sterilized, Non-Pyrogenic, Avantor). The 96-well plate was then incubated over the course of 72 hours at 37°C under anaerobic conditions (GasPak EZ Anaerobe Gas Generating Pouch System with Indicator, BD) with OD_600_ readings taken periodically over 72 hours with a microplate reader (Varikoskan LUX 3020-1160). The time spent to read the OD_600_ was minimized to ensure little disturbance and the GasPak was replaced at each time point. Growth curves were performed for each strain in each condition (maltotriose or glycogen). Data was fitted to a logistic growth model using GraphPad Prism version 10.1.0 for macOS (GraphPad Software, Boston, USA).

### Genomic DNA Extraction

Genomic DNA was extracted from 48-hour broth cultures (1.5 mL) of *G. swidsinskii* using a previously described modified salting out procedure (23). DNA concentration and A_260_/_A280_ ratio were measured using Nanodrop 2000c.

### RNA isolation and first strand cDNA synthesis

*Gardnerella swidsinskii* grown in individual wells of a 96 well plate (200 µL per well) was harvested 14 hours after incubation with OD_600_ values ranging from 0.3-0.6 (exponential growth phase) and pooled for a total of 800 µL of culture per sample. Cells were pelleted by centrifugation at 4°C and the supernatant was removed at which point they were immediately stored at -80°C until RNA isolation. The PureLink RNA Mini Kit (Invitrogen) and “Purifying RNA from Bacterial Cells” protocol was used with the following modifications. Mutanolysin and proteinase K were added to the lysozyme solution recommended by the manufactuer and a 15-minute incubation at 37°C after adding the lysozyme solution and 1% SDS was added. Once 350 µL of Lysis Buffer was added to each sample, two samples of the same strain and condition (glycogen or maltotriose) were pooled into one nuclease-free microcentrifuge tube therefore using double the recommended amount of lysozyme solution (total 200 µL) and Lysis Buffer (total 700 µL) per sample. The homogenization step was performed using syringes attached to 25G x 5/8” gauge needles and the optional on-column PureLink DNase Treatment was included. The manufacturer’s protocol was followed up to the elution step where the column was incubated for 2 minutes at room temperature with 30 µL of RNase free water before eluting. RNA concentration and A_260_/_A280_ were determined by spectrophotometry (Nanodrop 2000) and the RNA Integrity Number and concentration were measured using an Agilent 4200 TapeStation with RNA ScreenTape (5067-5576, Agilent) according to the manufacturer’s recommendations. A detailed laboratory protocol for the optimized RNA extraction is published elsewhere (24).

First strand cDNA was synthesized using Maxima First Strand cDNA Synthesis Kit for RT-qPCR with dsDNase (ThermoFisher Scientific). Template RNA (50 ng) was used, and a 30-minute incubation at 37°C for the dsDNase treatment. A 1:10 dilution of cDNA was performed in preparation of qPCR. Diluted cDNA was aliquoted into 50 µL and stored at -80°C.

### Primer design, validation, and optimization

Ten candidate reference genes were selected from the core genome of *Gardnerella swidsinskii* strains NR016, NR020, and NR021 ensuring they were conserved amongst all strains. We selected candidates representing a variety of functional categories (clusters of orthologous group (COG) categories). Primers were designed for each candidate reference gene and the two genes of interest using Primer3Plus (25). Validation of product specific amplification was performed by a conventional PCR assay with a thermal gradient (57.7°C to 71.6°C) with 20 ng genomic DNA. Each reaction was 25 µL containing 1× PCR buffer, 2.5mM MgCl_2_, 0.2mM dNTPs, 0.4 µM forward primer, 0.4 µM reverse primer, 0.05 U Taq DNA Polymerase (AccuStart II Taq DNA Polymerase, Avantor) and 20 ng of genomic DNA. The thermocycling parameters began with an initial denaturation at 95°C for 3 minutes followed by 40 cycles of (95°C for 30 seconds, T_anneal_ for 30 seconds and 72°C for 15 seconds) and a final extension at 72°C for 5 minutes. The products were visualized by agarose gel electrophoresis (1.5% w/v agarose) to select the best annealing temperature. Amplification efficiency for each primer set was calculated using real-time SYBR green PCR and a 10-fold dilution series of genomic DNA.

### qPCR

Each qPCR reaction was 10 µL containing 1× PowerUp SYBR Green Master Mix (A25742, ThermoFisher Scientific), 0.4 µM forward primer, 0.4 µM reverse primer and 2 µL template cDNA. Aliquots of cDNA were only thawed once to set up PCR. qPCR reactions were prepared using the QIAgility robotic pipetting system. The following amplification cycles were applied in a Bio-Rad CFX Connect real-time system: 1 cycle at 50°C for 2 minutes, 1 cycle at 95°C for 2 minutes, 40 cycles of 95°C for 15 seconds and 60°C for 15 seconds, followed by a melt curve, increasing the temperature by 0.5°C increments from 65°C to 95°C. Reaction mixture with no template DNA was used as a negative control and each biological replicate of *G. swidsinskii* in each condition had three technical replicates run in duplicate PCR reactions. The no-RT control was tested using the *ftsK* PCR.

### Gene expression stability and statistical analysis

Candidate reference gene expression stability was evaluated by analyzing the raw Cq values from the qPCR data with RefFinder (26) which combines four algorithms: the standard deviations of the delta-Cq (27), BestKeeper (28), NormFinder (29), and geNorm (11).

Relative gene expression was calculated using the formula [Relative Gene Expression = (E_GOI_) ^ΔCt^ ^GOI^/GeoMean[(E_REF_) ^ΔCq^ ^REF^]. Relative gene expression values were log_10_ transformed and statistical analysis was performed using an unpaired parametric t-test in GraphPad Prism version 10.1.0 for macOS (GraphPad Software, Boston, USA).

## Experimental design

*Gardnerella swidsinskii* strains NR016, NR020, and NR021 were grown on Columbia sheep blood agar plates anaerobically at 37°C for 48 hours from isolates stored at -80°C. Multiple colonies were then used to inoculate 5 mL mNYC III media supplemented with 1% (w/v) D-glucose and incubated anaerobically at 37°C for 48 hours. Aliquots (200 µL) of the diluted culture of each strain were then pipetted into separate flat bottom 96-well plates with 44 replicate wells per condition (maltotriose or glycogen). Uninoculated mNYC III media was used as a negative control with four replicates per condition. The 96-well plates were then incubated anaerobically at 37 °C until the mid-exponential phase (OD_600_ 0.3-0.6). Bacteria were harvested by pooling two wells (total volume of 400 µL) into pre-labelled microcentrifuge tubes and kept on ice while completing harvesting all bacteria. Bacteria were pelleted by centrifugation at 4°C, the supernatant was removed, and pellets were immediately stored at -80°C until RNA isolation. RNA extraction was performed in triplicate per strain (NR016, NR020, NR021) per condition (maltotriose, glycogen) using the PureLink RNA Mini Kit with modifications described in the Methods. An extraction control containing only kit reagents was included with each batch of extractions. RNA quality (A_260_/A_280_ and RIN) and quantity was examined using the Agilent TapeStation and Nanodrop 2000c. RNA extracts were stored in -80°C overnight before first strand cDNA synthesis using 50 ng of template RNA. cDNA synthesis was performed with the ThermoScientific Maxima First Strand cDNA Synthesis Kit for RT-qPCR with dsDNase, and a reverse transcriptase minus (RT-) negative control and no template control (NTC) were included for each set of extractions. qPCR was performed with two technical replicates for each cDNA sample. A total of 36 Cq values were obtained for each candidate reference gene and gene of interest (3 isolates × 2 conditions × 3 RNA/cDNA preps × 2 PCR replicates).

## Results

### Growth curves and RNA purification

Growth curves were measured to determine the relationship between OD_600_ and the various phases of growth, and mid-exponential phase corresponded to an OD_600_ value of 0.3-0.6 for all strains of *Gardnerella swidsinskii* incubated in a mNYC III medium supplemented with 1% maltotriose or glycogen (Figure S1). No growth was observed in the media only controls.

RNA was purified from cultures grown in 96-well plates according to the experimental design described in the Methods. A total of 18 extractions were performed where three RNA extractions were performed per isolate (NR016, NR020 and NR021) in each condition (maltotriose and glycogen). RNA extracts contained 1.1 - 4.1 µg of RNA (mean 2.3 µg), with A_260_/A_280_ ratios of 2.06 - 2.24 (median 2.13) and RIN values 7.5 - 8.9 (median 8.05) (Table S1).

### Primer validation

Primers were evaluated using genomic DNA to assess specific amplification of products and efficiency (Table 1). Primers targeting eight of our ten candidate reference genes and both of our genes of interest specifically amplified their intended target (Figure S2). Primers for *fusA* and *glnA1* did not specifically amplify their targets and therefore were not further evaluated. Amplification efficiencies were ranged from 85.30% to 98.20% (Table 1) with all except the *pgm* assay being >90% efficient. Observation of single melt peaks following qPCR further confirmed primer specificity and lack of primer dimers for all candidate reference genes (Figure S3).

**Table 1.**
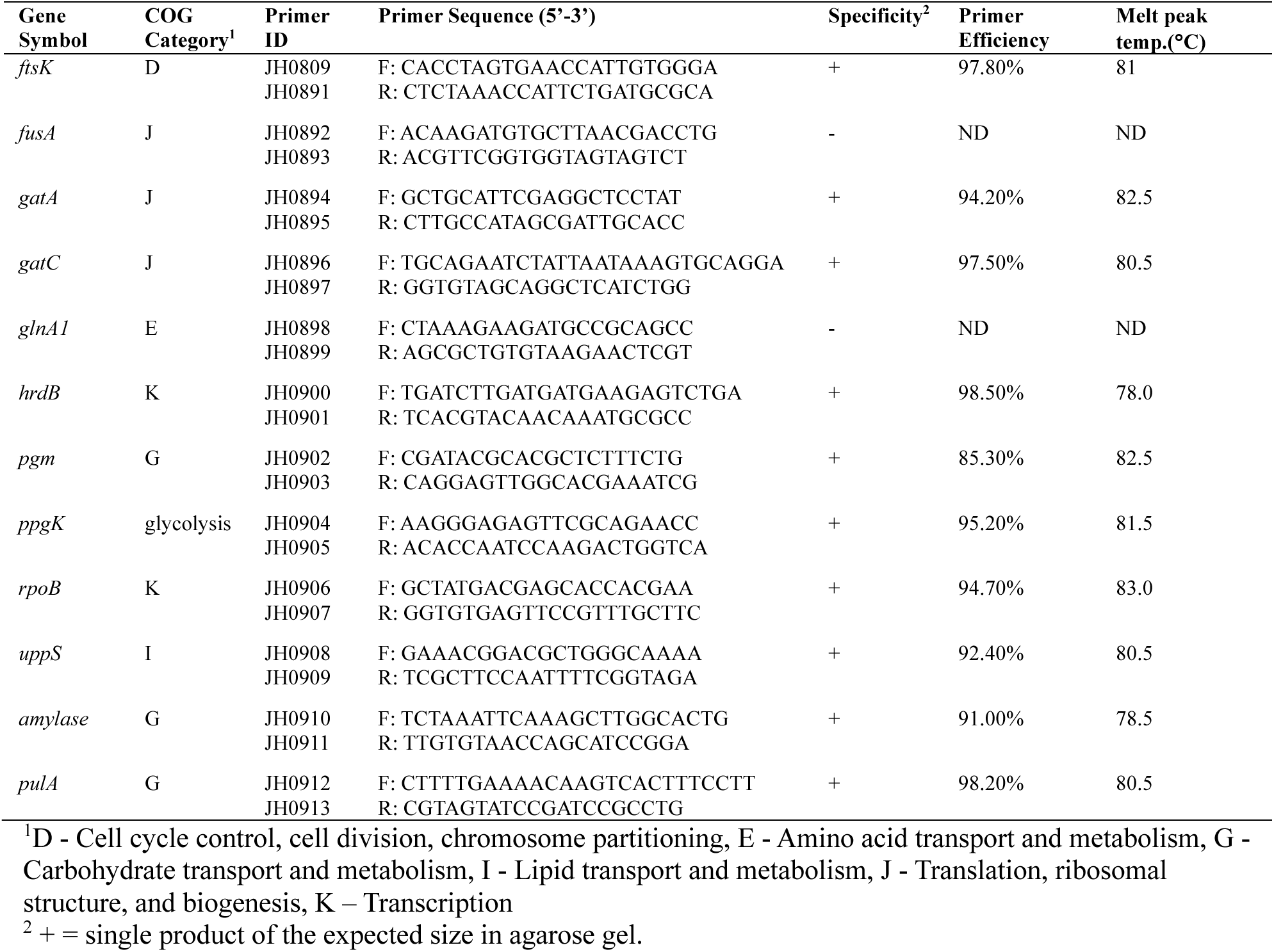
Primer sequences, clusters of orthologous groups (COGs), and results of primer validation testing specificity, efficiency, and melt temperatures of PCR products of candidate reference genes and genes of interest evaluated in this study.

### Evaluation of stability of candidate reference genes

The expression stability of candidate reference genes (*ftsK, gatA, gatC, hrdB, pgm, ppgK, rpoB* and *uppS*) was assessed in media supplemented with maltotriose or glycogen by RT-qPCR. Cq values for this experiment ranged from 15.63 to 22.92 (Figure 1). Candidate reference gene expression stability was assessed using RefFinder which combines four algorithms: Delta-Cq, BestKeeper, NormFinder, geNorm.

**Figure 1.**
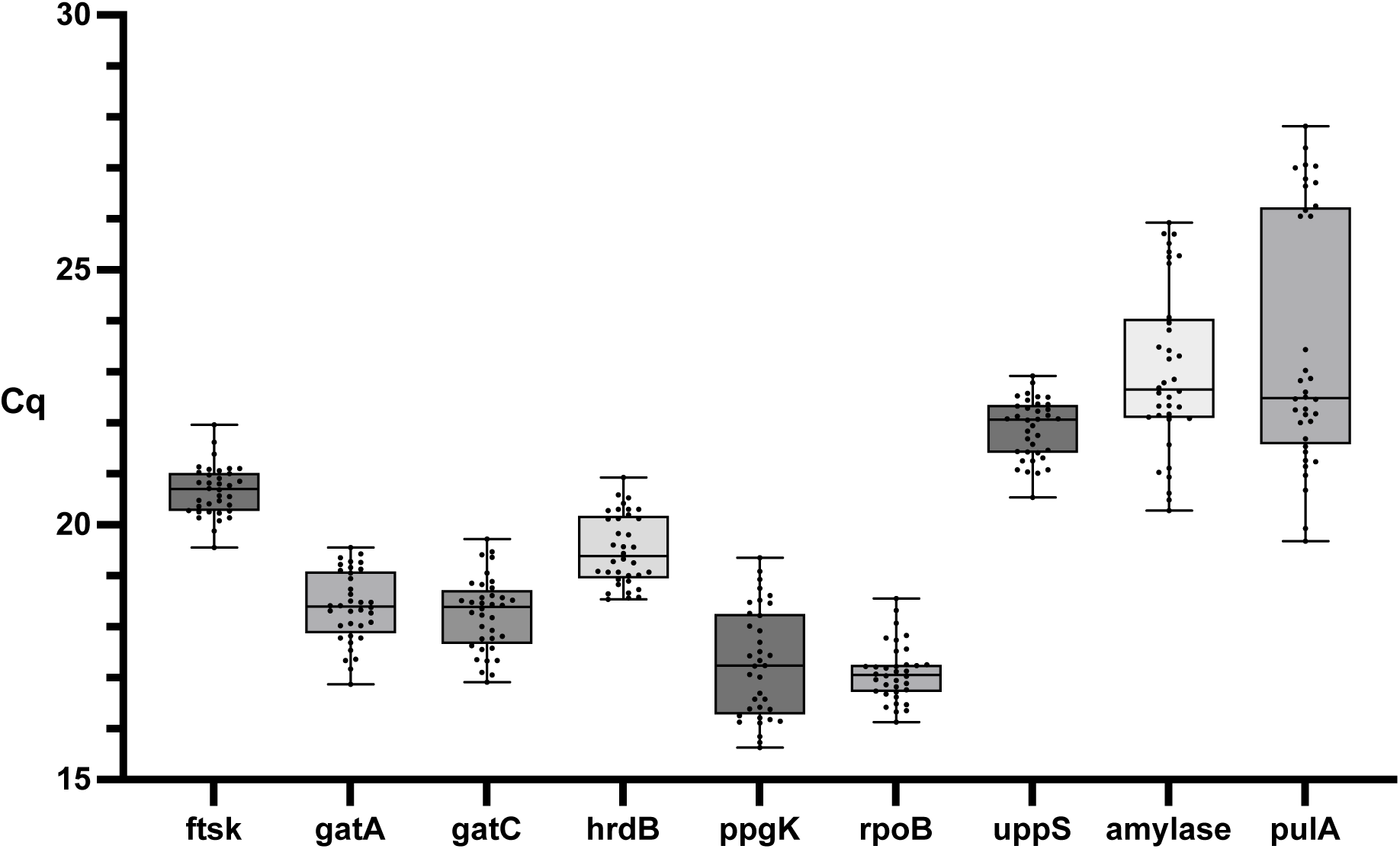
Distribution of Cq values of candidate reference genes across all samples. The boxplot communicates the 25^th^ and 75^th^ percentile represented by the lower and upper box boundaries respectively, the median represented by the line inside the box, and the minimum and maximum represented by the error lines. Each boxplot constitutes 36 Cq values which are plotted as individual points (3 isolates, 2 conditions, 3 RNA/cDNA preps, 2 technical PCR replicates).

#### Delta-Cq

The mean of the standard deviation derived from a comparison between a reference gene and any other candidate is calculated as the gene stability indicator where a lower arithmetic mean indicates higher stability of the gene. In ranking order, *uppS* had the lowest arithmetic mean followed by *gatA, gatC, rpoB, ftsK, hrdB, ppgK*: 0.70, 0.72, 0.75, 0.79, 0.84, 0.87, 1.05.

#### BestKeeper

Correlations of the expression levels of all candidate reference genes are estimated and highly correlated ones are combined into an index. From this, the standard deviation, percent covariance, and power of the candidates are calculated. Reference genes whose standard deviation is below 1.0 are stable and the lower the standard deviation, the higher the stability. *ftsK* had the lowest standard deviation followed by *rpoB, uppS, gatA, gatC, hrdB, ppgK*: 0.395, 0.419, 0.498, 0.556, 0.556, 0.584, 0.6, 0.909. All candidate reference genes fell below the threshold of 1.0 indicating stability according to BestKeeper.

#### NormFinder

NormFinder not only estimates the overall variation of the candidate reference genes but also the variation between subgroups of samples. NormFinder allows for a direct measurement for the estimated expression variation. *uppS* had the lowest variation followed by *gatA, gatC, rpoB, ftsK, hrdB, ppgK*: 0.350, 0.411, 0.470, 0.546, 0.626, 0.653, 0.917.

#### geNorm

By calculating the average pairwise variation of a particular gene with all other genes, all candidate reference genes are ranked based on average expression stability value (M value). Stepwise exclusion of the least stable gene (highest M value) is performed until the two most stable genes cannot be further ranked. Genes below a threshold of 1.5 are stable. The two most stable genes according to the M value were *gatA* and *gatC*, followed by *uppS, rpoB, ftsK, hrdB, ppgK*: 0.494, 0.538, 0.622, 0.676, 0.725, 0.818. All candidate reference genes fall below the 1.5 threshold.

#### RefFinder

A comprehensive ranking by RefFinder is provided by calculating the geometric mean of the rankings of all candidate reference genes by the four algorithms. Based on this ranking, *uppS* was the top comprehensively ranked reference gene followed by *gatA, gatC, ftsK, rpoB, hrdB, ppgK* with geometric ranking values of 1.73, 2.00, 2.59, 3.34, 3.36, 6.00, 7.00, respectively (Figure 2).

**Figure 2.**
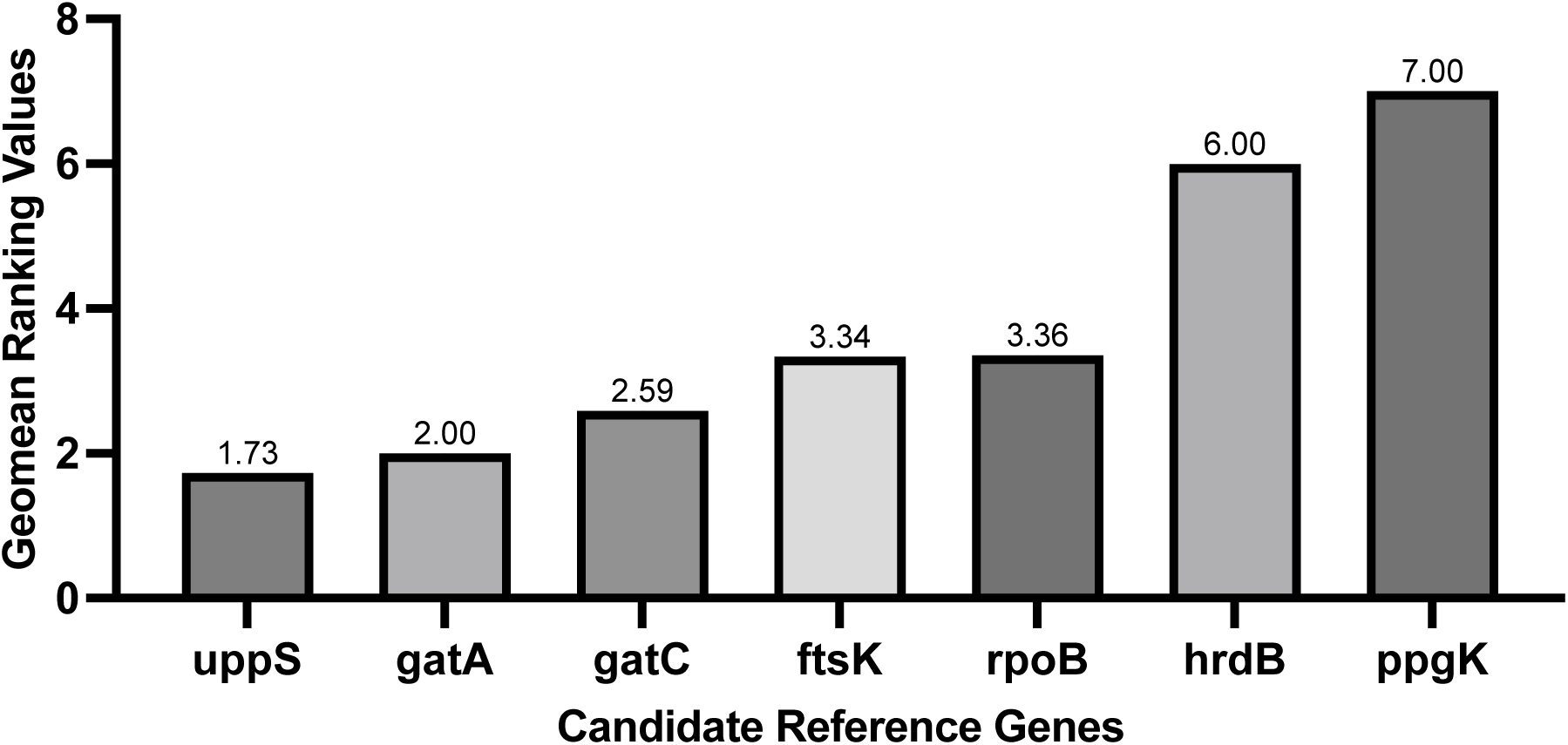
RefFinder ranking of candidate reference genes. Seven candidate reference genes were comprehensively ranked by calculation of the geometric mean of rankings based on delta-Cq, BestKeeper, NormFinder and geNorm.

### α-amylase and α-amylase-pullulanase relative gene expression

As outlined in the experimental design presented in the Methods, the expression of α-amylase and α-amylase-pullulanase was assessed in media supplemented with 1% glycogen or 1% maltotriose by RT-qPCR using the cDNA synthesized from the RNA extractions described in Table S1. Cq values for α-amylase and α-amylase-pullulanase, regardless of strain or condition, were 20.28-25.93 and 19.68-27.82, respectively. When strain and condition were considered, we observed that strain NR021 in both glycogen and maltotriose had higher Cq values for both genes compared to the NR016 and NR020 strain (Figure 3).

**Figure 3.**
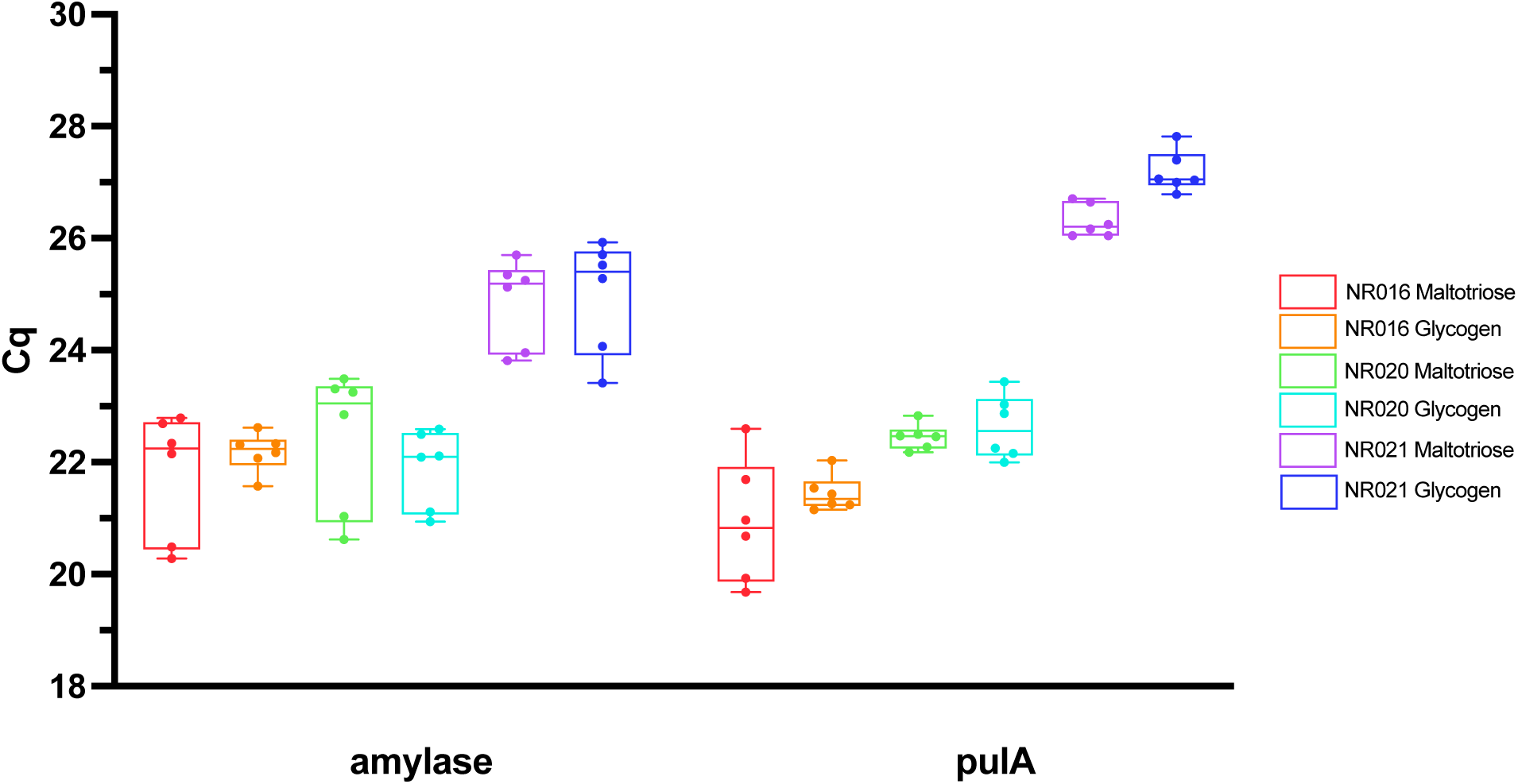
Distribution of Cq values for genes encoding α-amylase and α-amylase-pullulanase across all samples. The boxplot shows the 25^th^ and 75^th^ percentile represented by the lower and upper box boundaries, respectively, the median is represented by the line inside the box, and the minimum and maximum are represented by the whiskers. Each boxplot represents six Cq values, which are also shown as individual points.

We took advantage of our identification of multiple stable reference genes and applied the relative gene expression equation [Relative Gene Expression =(E_GOI_) ^ΔCt^ ^GOI^/GeoMean[(E_REF_) ^ΔCq^ ^REF^] to determine relative expression of our genes of interest in the two growth conditions. This approach incorporates multiple reference genes and so we selected the top two comprehensively ranked reference genes, *uppS* and *gatA*. The relative gene expression of α-amylase ranged from 0.40-1.64 in media supplemented with maltotriose and 0.40-2.29 in media supplemented with glycogen. The relative gene expression of α-amylase-pullulanase ranged from 0.39-2.16 in media supplemented with maltotriose and 1.18-2.66 in media supplemented with glycogen. To facilitate comparison of ratios, relative expression values were log_10_ transformed and compared using an unpaired parametric t-test. Log_10_ relative gene expression of α-amylase did not differ significantly in the two growth conditions (p = 0.4766) while log_10_ relative gene expression of α-amylase-pullulanase was significantly higher in bacteria grown in glycogen supplemented media compared to media supplemented with maltotriose (p = 0.0064) (Figure 4).

**Figure 4.**
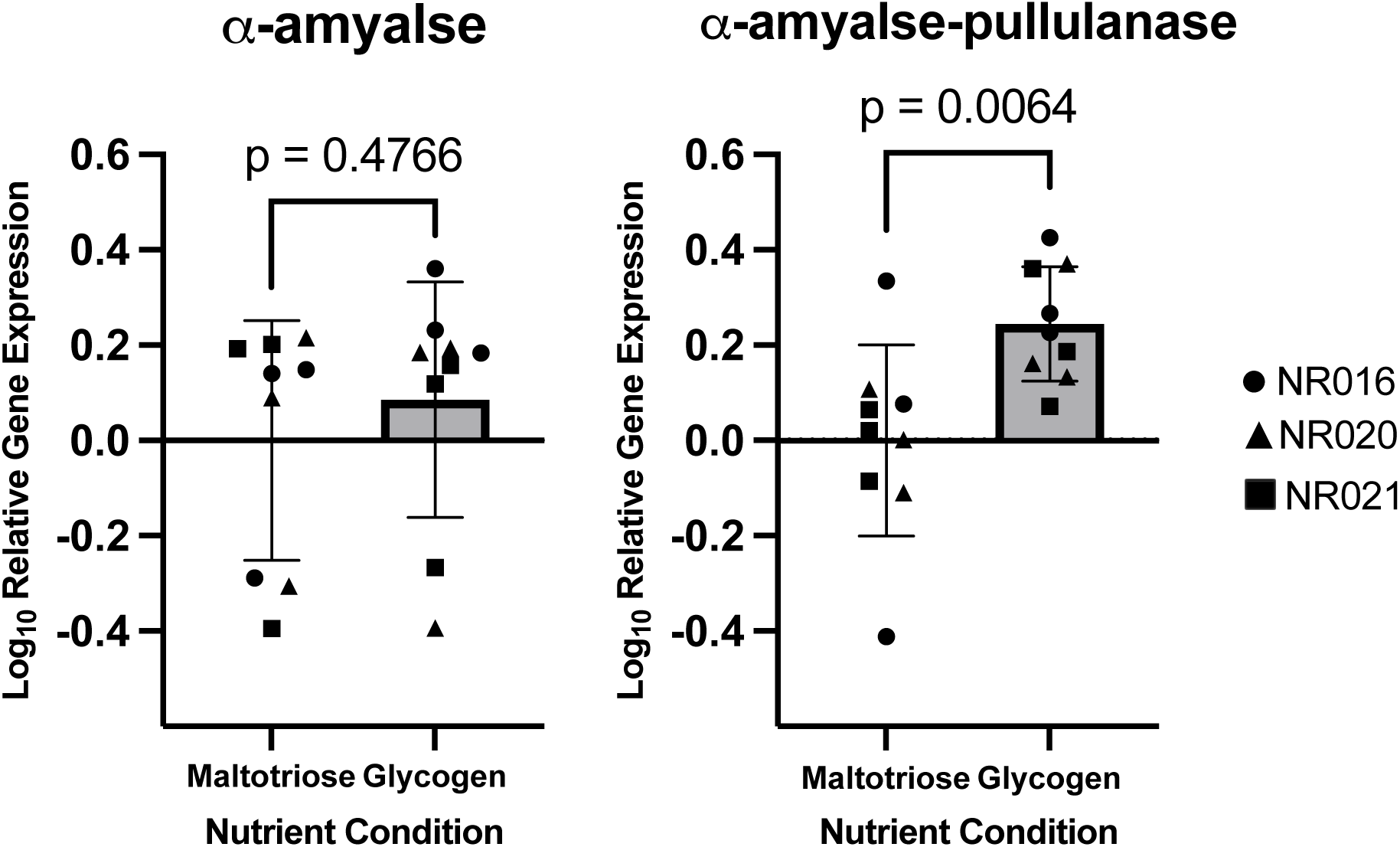
Log_10_ relative gene expression of α-amylase (left) and α-amylase-pullulanase (right) in mNYC III media supplemented with 1% maltotriose or glycogen. Error bars indicate standard deviation. Statistical analysis was performed using an unpaired parametric t-test with p < 0.05 considered significant.

## Discussion

The overgrowth of *Gardnerella* spp. within the human vagina is an important clinical indication of bacterial vaginosis (30). Development of improved diagnostics and clinical interventions for BV will be facilitated by improved understanding of the growth and colonization of these bacteria in relation to utilization of available nutrients in the vagina. Glycogen is one of the most important and abundant nutrient sources in the vaginal lumen that is degraded by α-amylase and α-amylase-pullulanase into smaller malto-oligosaccharides utilized by the vaginal microbiota (6, 9). Differential expression of genes encoding glycogen degrading enzymes in response to relative abundance of relevant substrates and products has not previously been investigated for *Gardnerella* spp. To conduct gene expression studies of these enzymes, reference genes whose gene expression is unaffected by the conditions of the study, must be identified.

Accurate normalization of data is necessary to avoid erroneous differences in target gene expression (31–33). In this study, we identified suitable reference genes for *Gardnerella swidsinskii* in media supplemented with 1% maltotriose or glycogen. RefFinder found *uppS* to be the top comprehensively ranked reference gene followed by *gatA*, *gatC*, *ftsK*, *rpoB*, *hrdB*, and *ppgK*. All candidate reference genes fell below the threshold determining gene stability according to geNorm and BestKeeper. The primers designed for reference gene PCR assays are specific for *G. swidsinskii*, so although these genes may also be appropriate for used in gene expression studies in other species of *Gardnerella*, modifications would be needed to make the assays optimal for other species.

With the identified reference genes, targeted gene expression analysis of the glycogen degrading enzymes, α-amylase and α-amylase-pullulanase, was possible. The delta-delta Cq method is commonly applied in calculating relative gene expression from RT-qPCR experiments (34), however, this method is limited in accounting for differences in primer efficiencies and does not allow for the use of more than one reference gene for analysis. Therefore, we applied a variation of the Pfaffl method which overcomes these limitations (35). While no significant difference in log_10_ relative gene expression of α-amylase in the media supplemented with 1% maltotriose or glycogen was observed, log_10_ relative gene expression α-amylase-pullulanase was significantly higher in media supplemented with 1% glycogen relative to media supplemented with 1% maltotriose (Figure 4).

Glycogen digestion is achieved by the collective action of “amylase” enzymes, which cleave α-1,4 and α-1,6 glycosidic bonds either from the non-reducing end or randomly within the chain. The single domain α-amylase enzyme of *G. swidsinskii* hydrolyzes α-1,4 glycosidic bonds producing maltose, maltotriose, and maltotetraose from glycogen (9). The α-amylase-pullulanase enzyme contains two catalytic domains: the α-amylase and pullulanase domains (9). The α-amylase domain has similar activity to the α-amylase single domain enzyme, while the pullulanase domain contributes additional de-branching capabilities, acting on α-1,6 glycosidic bonds. We have shown previously that malto-oligosaccharides including maltotriose are preferred growth substrates for *Gardnerella* spp. (10). Maltotriose contains only α-1,4 linkages and lacks any branching. Our observation of increased expression of α-amylase-pullulanase but not α-amylase in media supplemented with 1% glycogen relative to media supplemented with 1% maltotriose may be explained by the functional differences in these two enzymes. In media rich with glycogen and therefore α-1,6 glycosidic bonds, α-amylase-pullulanase expression may be upregulated to digest these bonds, however, when maltotriose is abundant, there would be no need for the debranching activity of α-amylase-pullulanase. Production and export of the large (>100 kDa) α-amylase-pullulanase enzyme would come with a fitness cost that might not be worthwhile when preferred nutrients are abundant.

Carbon catabolite repression is a regulatory mechanism in some bacteria by which the expression of genes essential for the utilization of secondary carbon sources is prevented by the presence of a preferred substrate (36). This phenomenon has been observed in some *Lactobacillus* spp., including vaginal *L. crispatus* (37). In these bacteria, the pullulanase-encoding pulA gene promoter contains a binding site for CcpA (carbon catabolite protein A), a global regulator of carbon metabolism in Gram positive bacteria that has been shown to be involved in repressing alternative carbohydrate utilization pathways in the presence of abundant glucose (38–40). Glucose is not a preferred substrate for *Gardnerella* spp. (10) and whether *G. swidsinskii* possess a carbon catabolite repression mechanism that regulates expression of glycogen degrading enzymes will be explored in future studies.

Despite its clinical significance and long-standing association with the most common vaginal condition of reproductive aged women, knowledge of many of the basic biological attributes of *Gardnerella* spp. are undescribed. Both genome-wide transcriptome studies and targeted gene expression studies using RT-qPCR aimed at determining how *G. swidsinskii* reacts and adapts to changes in its external environment will be facilitated by the protocol for high quality RNA isolation and the identification of reference genes for *G. swidsinskii* presented here. Our results also present tools that are directly applicable to gene expression studies in other *Gardnerella* spp.

## Acknowledgements

This work was supported by a Natural Sciences and Engineering Research Council of Canada (NSERC) Undergraduate Student Research Award (USRA). The authors are grateful to Dr.

Maarten Voordouw for helpful discussions on statistical analysis.

## Supplemental Material

**Table S1.** RNA yield and quality examined via Agilent TapeStation (RIN values) and spectrophotometry (A_260_/A_280_) for each RNA extraction performed.

| Sample ID | RNA Yield<br>(ng/μL) | RNA Integrity Number<br>(RIN) | $A_{260}/A_{280}$ |
| --- | --- | --- | --- |
| NR016 Maltotriose Replicate 1 | 46.9 | 7.7 | 2.24 |
| NR016 Maltotriose Replicate 2 | 85.9 | 8.1 | 2.15 |
| NR016 Maltotriose Replicate 3 | 77.0 | 8.0 | 2.12 |
| NR016 Glycogen Replicate 1 | 49.3 | 7.5 | 2.16 |
| NR016 Glycogen Replicate 2 | 58.5 | 7.9 | 2.14 |
| NR016 Glycogen Replicate 3 | 37.4 | 7.5 | 2.14 |
| NR020 Maltotriose Replicate 1 | 92.0 | 8.9 | 2.14 |
| NR020 Maltotriose Replicate 2 | 72.2 | 8.7 | 2.14 |
| NR020 Maltotriose Replicate 3 | 136.0 | 8.8 | 2.13 |
| NR020 Glycogen Replicate 1 | 99.0 | 8.9 | 2.17 |
| NR020 Glycogen Replicate 2 | 69.3 | 8.7 | 2.12 |
| NR020 Glycogen Replicate 3 | 75.9 | 8.8 | 2.07 |
| NR021 Maltotriose Replicate 1 | 132 | 8.1 | 2.11 |
| NR021 Maltotriose Replicate 2 | 67.0 | 8.0 | 2.06 |
| NR021 Maltotriose Replicate 3 | 69.2 | 7.8 | 2.11 |
| NR021 Glycogen Replicate 1 | 56.2 | 7.9 | 2.10 |
| NR021 Glycogen Replicate 2 | 51.2 | 7.9 | 2.14 |
| NR021 Glycogen Replicate 3 | 94.5 | 8.3 | 2.08 |

**Figure S1.**
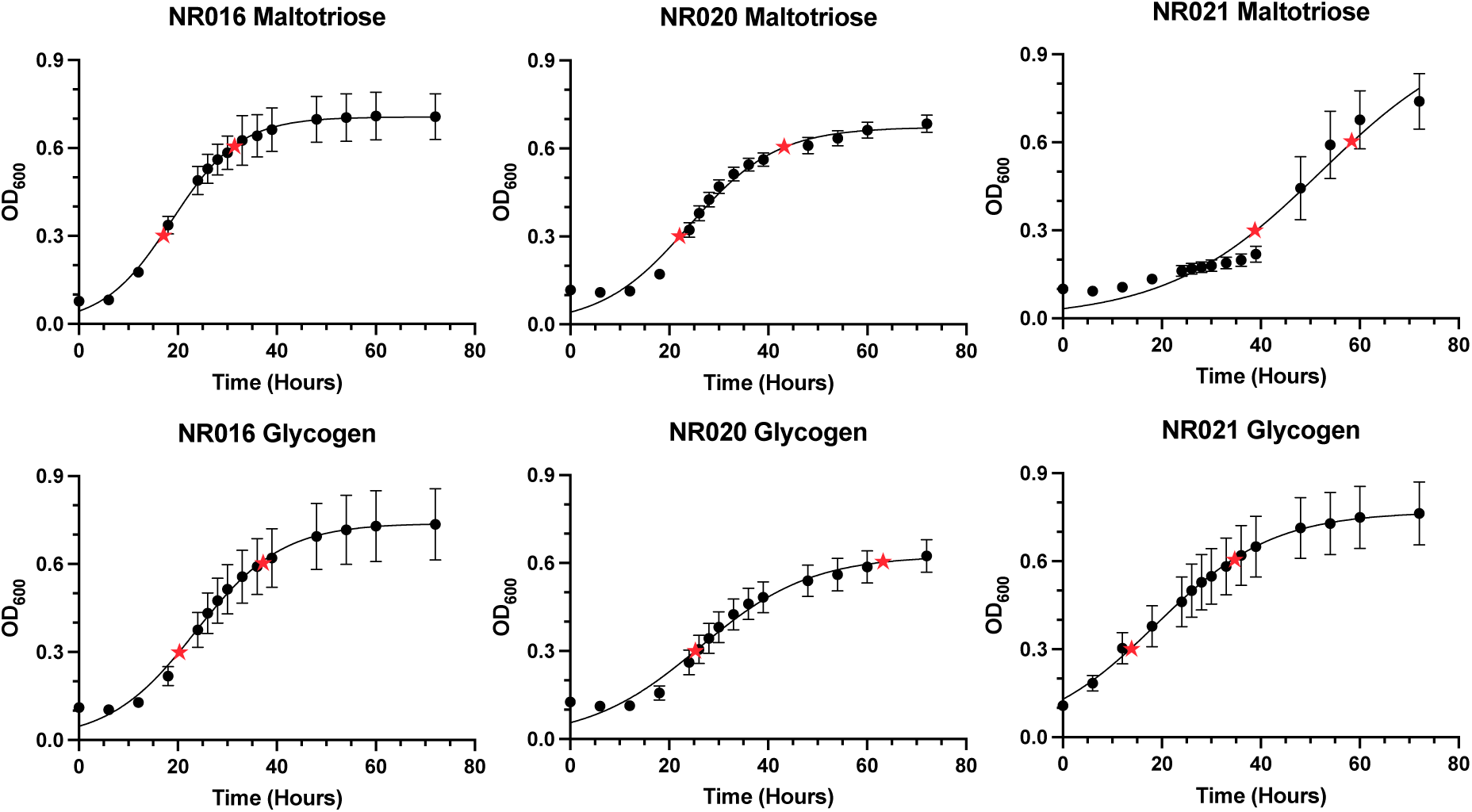
Growth curves over 72 hours of *Gardnerella swidsinskii* strains NR016, NR020, and NR021 grown in a mNYC III medium supplemented with 10% heat inactive bovine serum and 1% glycogen or maltotriose. Data represents mean ± SD of eight replicates. Mid-exponential phase was determined to be at an OD_600_ reading with the microplate reader at 0.3-0.6 marked by the red stars.

**Figure S2.**
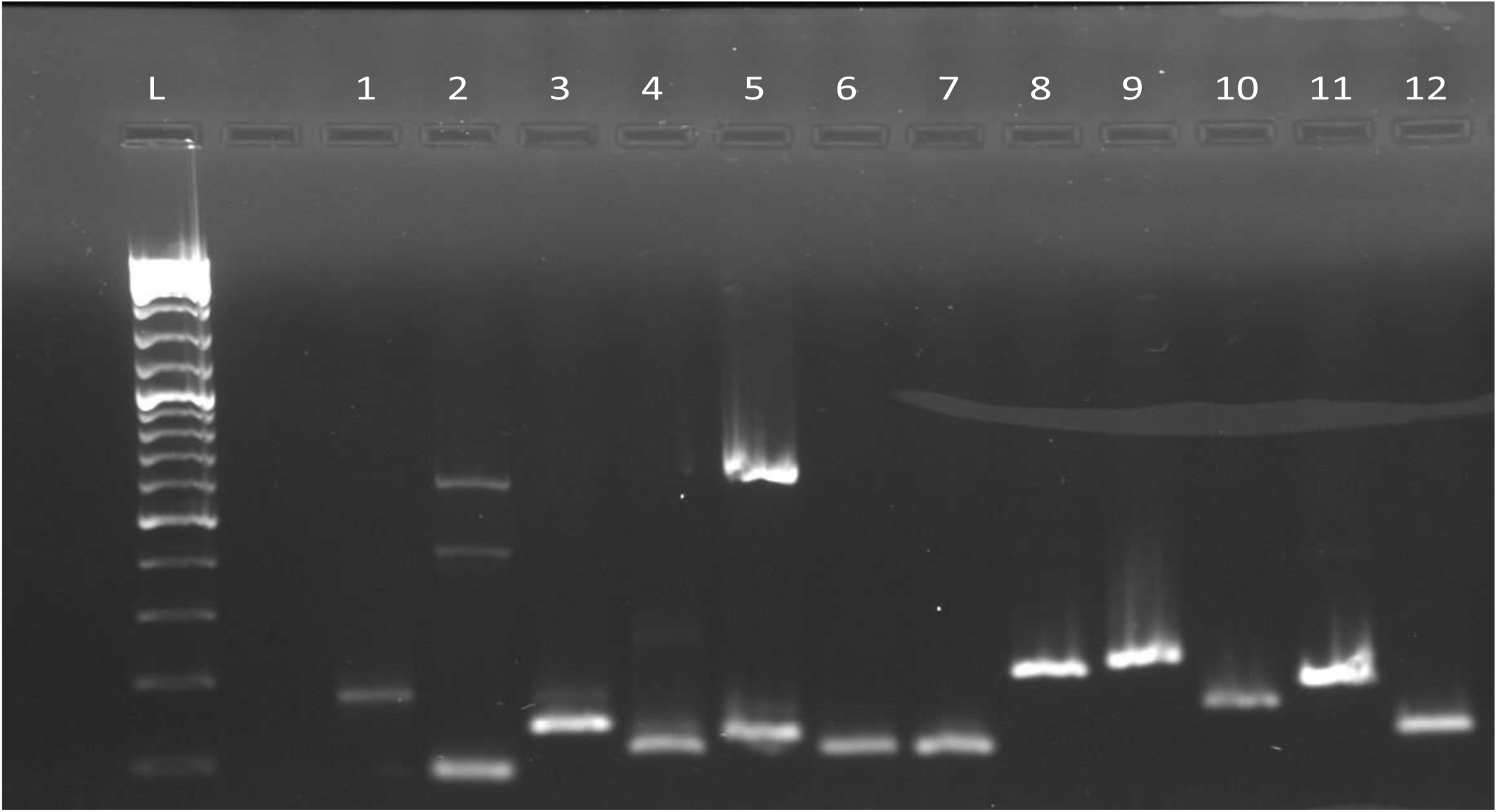
1.5% agarose gel electrophoresis, 100V for 40 minutes, 0.5X TBE of PCR amplified products using genomic DNA of NR021 *G. swidsinskii* with all primers (L-GeneRuler™ DNA Ladder Mix (SM0331, ThermoFisher Scientific), Lane 1-ftsK, 2-fusA, 3-gatA, 4-gatC, 5-glnA, 6-hrdB, 7-pgm, 8-ppgK, 9-rpoB, 10-uppS, 11-amylase, 12-pulA).

**Figure S3.**
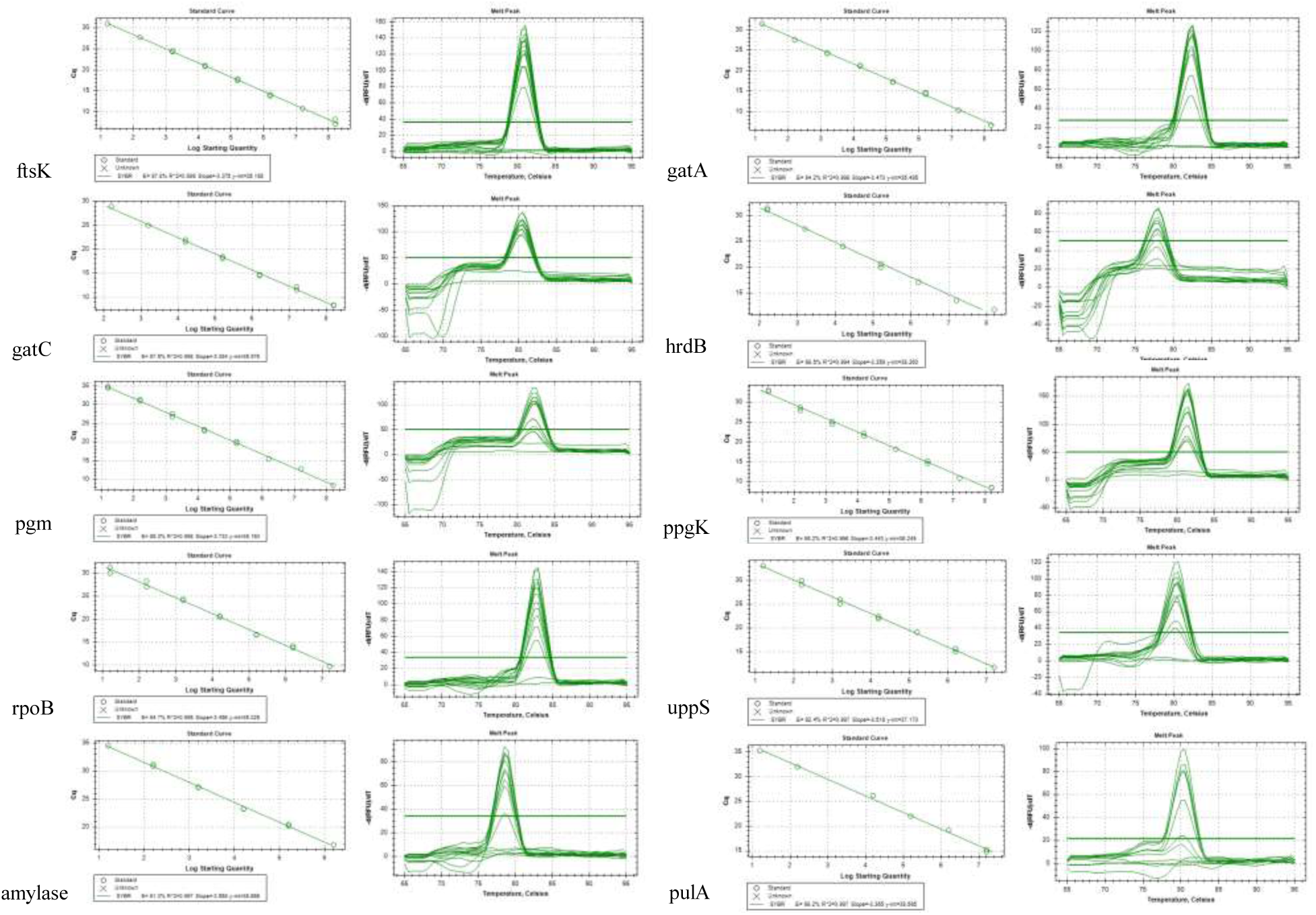
Standard curves and melt peaks constructed of select candidate reference genes and genes of interest.

